# Quantifying the potential causes of Neanderthal extinction: abrupt climate change versus competition and interbreeding

**DOI:** 10.1101/2020.04.19.049734

**Authors:** Axel Timmermann

**Affiliations:** Center for Climate Physics, Institute for Basic Science, Busan, South Korea; Pusan National University, Busan, South Korea

**Keywords:** Neanderthal extinction, Pleistocene, abrupt climate change, numerical modeling, Climate Dynamics

## Abstract

Anatomically Modern Humans are the sole survivor of a group of hominins that inhabited our planet during the last ice age and that included, among others, *Homo neanderthalensis*, *Homo denisova*, and *Homo erectus*. Whether previous hominin extinctions were triggered by external factors, such as abrupt climate change, volcanic eruptions or whether competition and interbreeding played major roles in their demise still remains unresolved. Here I present a spatially resolved numerical hominin dispersal model (HDM) with empirically constrained key parameters that simulates the migration and interaction of Anatomically Modern Humans and Neanderthals in the rapidly varying climatic environment of the last ice age. The model simulations document that rapid temperature and vegetation changes associated with Dansgaard-Oeschger events were not major drivers of global Neanderthal extinction between 50-35 thousand years ago, but played important roles regionally, in particular over northern Europe. According to a series of parameter sensitivity experiments conducted with the HDM, a realistic extinction of the Neanderthal population can only be simulated when *Homo sapiens* is chosen to be considerably more effective in exploiting scarce glacial food resources as compared to Neanderthals.

## 1. Introduction

The rapid demise of Neanderthals (Higham et al., 2014; Mellars, 2004) occurred ∼50-35 ka (Benazzi et al., 2011; Higham et al., 2011) - shortly after Anatomically Modern Humans (AMHs) dispersed into subtropical and extratropical Eurasia. A variety of hypotheses have been put forward to explain the extinction of Neanderthals, including competitive exclusion (Banks et al., 2008; Flores, 1998), assimilation (Smith et al., 2005), demographic weakness (Degioanni et al., 2019), abrupt climate and vegetation change (Finlayson and Carrion, 2007; Staubwasser et al., 2018), pathogens (Houldcroft and Underdown, 2016), and volcanic eruptions (Fitzsimmons et al., 2013). The proposed mechanisms are not mutually exclusive, and their relative role may have also varied regionally and temporally.

The goal of this study is to quantify the importance of resource competition, climate change and hybridization in the large-scale Neanderthal extinction across Eurasia using a 2-dimensional numerical hominin dispersal model driven by estimates of past climate changes and by conducting a series of targeted sensitivity experiments. Previous idealized numerical studies on the disappearance of Neanderthals relied mostly on zero-dimensional non-spatial models (Gilpin et al., 2016; Roberts and Bricher, 2018) and often ignored the effects of realistic glacial climate forcing.

The key premises of my modeling approach are that i) migration, growth, mortality, competition and interbreeding of AMH and Neanderthals can be expressed by a set of coupled nonlinear Fisher-Kolmogorov reaction-diffusion equations (Gilpin et al., 2016; Timmermann and Friedrich, 2016) (**section 2**); ii) the effect of past climate change on hominin growth and dispersal can be described by changes in annual mean terrestrial net primary production (NPP) (Eriksson et al., 2012; Vandati et al., 2019), temperature and desert fraction; iii) northward migrating AMHs started to adapt to reduced sunlight (Beleza et al., 2013; Crawford et al., 2017) in subtropical and extratropical regions between 60-40 ka, which helped them expand into Eurasia; iv) compared to AMH, Neanderthals remained in a climatically and environmentally more confined region (Fabre et al., 2009), and they likely lived in smaller, more scattered groups as suggested by a number of studies (Castellano et al., 2014; Melchionna et al., 2018). Using these key assumptions, the Hominin Dispersal Model, version 2 (HDM2) presented here (see Methods section) simulates the spatio-temporal evolution of the density of *Homo sapiens* (HS) and *Homo neanderthalensis* (HN) in response to time-varying patterns of temperature, NPP, desert fraction and sea level (**Supplementary Movie 1,2**). The climatic information is obtained by combining data from a transient earth system model (Goosse et al., 2010), which has been forced for 784 ka (Friedrich et al., 2016) with orbitally-driven changes in solar insolation greenhouse gas concentrations and estimates of Northern Hemisphere ice-sheet extent (Ganopolski et al., 2010), with a semi-empirical estimate of millennial-scale variability (Figure 1).

**Figure 1.**
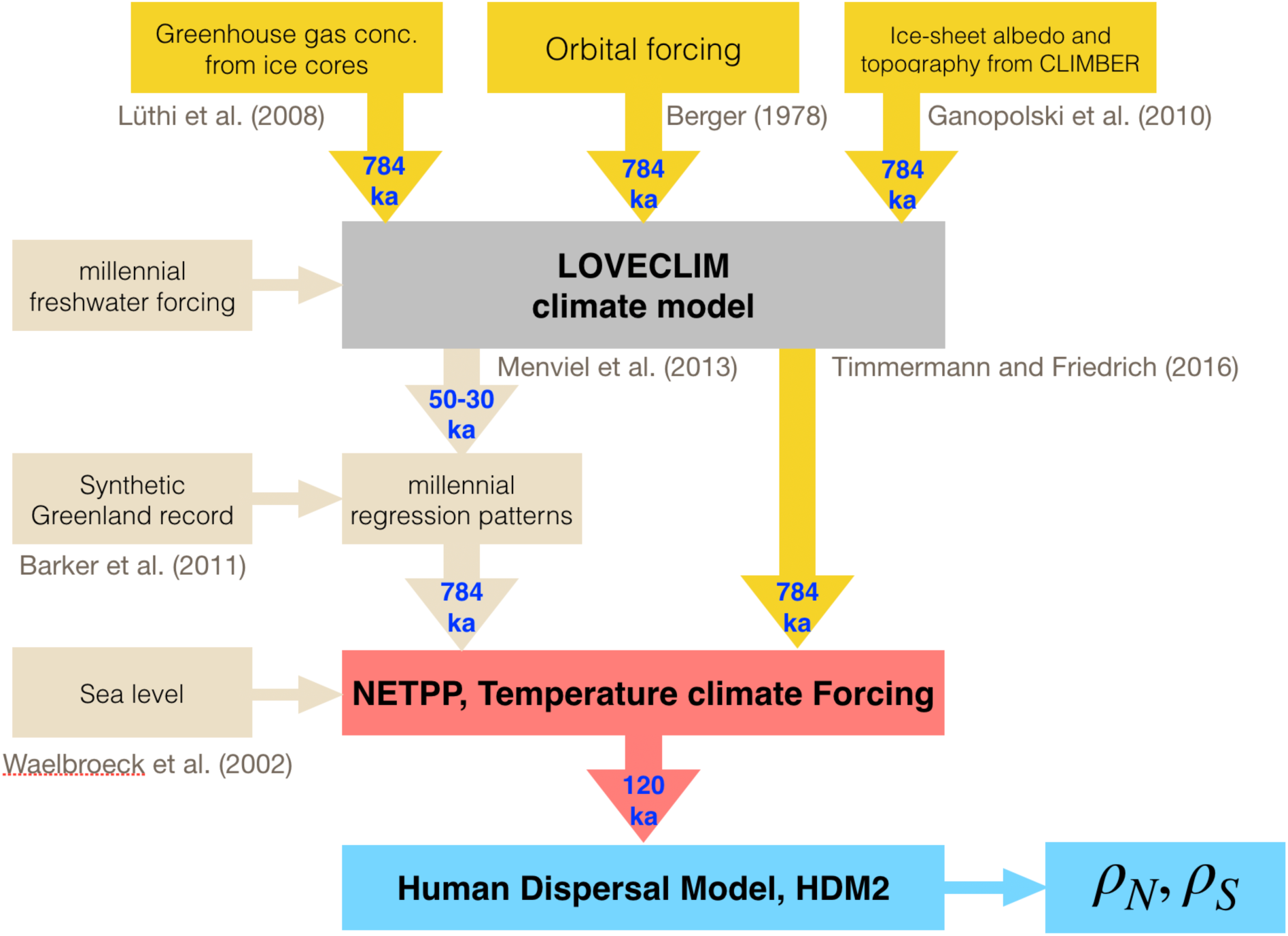
Schematics of climate forcing for Human Dispersal Model, version 2. The climate forcing (temperature, net primary production and desert fraction) for HDM2 are obtained by combining the direct simulated values from an orbital-scale transient LOVECLIM glacial-interglacial simulation (ORB) with a statistical estimate of millennial-scale variability, which combines paleo-climate timeseries and millennial-scale characteristic patterns for temperature, net primary production and desert fraction from another millennial-scale LOVECLIM simulation.

After introducing the modelling framework and the key experiments in section 2, I present the main results from the parameter sensitivity experiments (section 3). The discussion focuses on the effect of interbreeding, competition and Dansgaard-Oeschger variability on Neanderthal extinction. In section 4 the main results will be summarized and discussed in the context of genetics, archaeology and paleoanthropology.

## 2. Method and Experiments

### 2.1 Hominin Dispersal Model

To simulate the dispersal and interaction of hominins in a time-varying environment a modified Fisher-Kolmogorov model is adopted. It makes the assumption that the dispersal of individuals can be described by a continuous partial differential equation (PDE) for the phenomenological mean field population density per area. Compared to our previous study (Timmermann and Friedrich, 2016), I included a second prognostic equation to capture the temporal evolution of Neanderthal densities. I also updated the numerical scheme (Praprotnik et al., 2004) and the overall formulation of carrying capacity. A more complex representation of diffusivity is included, which parameterizes the effect of topographic gradients and rivers. I also applied an updated climate forcing, which represents the observed millennial-scale Dansgaard-Oeschger variability more accurately (Fig. 1b).

The HDM model (version 2, **HDM2**) is based on two coupled dynamical Reaction-Diffusion Equations, which capture the time-evolution of the density of HN and HS as a function of latitude (y) and longitude (x). The PDEs are formulated as:

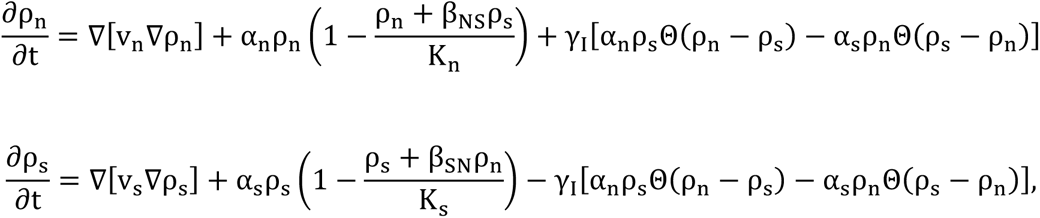

where ρ_n_(x,y) and ρ_s_(x,y) captures the density per area of *Homo Neanderthalensis* (HN) and *Homo sapiens* (HS) in each grid box with longitude x and latitude y, respectively. v_n_ and v_s_ denote the corresponding spatially dependent diffusivities and Θ(x) is the Heaviside function. The first terms describe the diffusion of hominins, whereas the other terms - referred to as reaction terms - represent the logistic growth process with α_n_, α_s_ representing the growth rates of Neanderthals and *Homo sapiens* and K_n_ and K_s_ the carrying capacity, respectively. Food competition between HN and HS is parameterized through the variables β_NS_, β_SN_. HDM2 simulates interbreeding between HS and HN. This parameter controls the effectiveness with which HS exploited existing food resources (represented through K_n_ and K_s_) relative to HN. It may capture a wide range of processes including differences in hunting strategies, social behavior, technological and physiological characteristics. It becomes evident that this parameter is highly uncertain, and it will there be chosen as a key parameter for an ensemble of sensitivity experiments. Here I assume that, if HS and HN populate the same grid cell, but in different densities, a certain fraction of the new births of the smaller group will come from interbreeding each time step with the larger group, generating offspring, that belongs to the larger group. The respective children are then removed from the smaller group and added to the larger group. This admixture process is represented by the last terms and the average interbreeding factor is abbreviated as γ_I_. Representing other interbreeding scenarios, in which for instance off-spring stays with the mother’s group, is beyond the scope of this paper. Such behavior can be simulated more appropriately using Individual Based Models (Vandati et al., 2019), which can include gender and age.

The carrying capacities K_n_(x,y,t), K_s_(x,y,t) of the logistic growth terms are calculated from the net primary production on land [N(x,y,t)], obtained by adding a simulated orbital-scale N_o_(x,y,t) contribution from the LOVECLIM transient forced 784 kyr simulation (Friedrich et al., 2016) and a reconstructed millennial-scale contribution N_m_(x,y.t), which is obtained by projecting a millennial-scale index timeseries(Barker et al., 2011) T_G_’(t) onto a characteristic millennial-scale regression pattern P_N_(x,y), described further below. One obtains N(x,y,t)=[N_o_ (x,y,t)+ T_G_’(t) P_N_(x,y)] Θ[N_o_ (x,y,t)+ T_G_’(t) P_N_(x,y)].

HN likely lived in smaller (Bocquet-Appel and Degioanni, 2013) and more scattered groups (Rogers et al., 2017), and had a smaller long-term effective population size as compared to HS and likely lower genetic diversity (Castellano et al., 2014). Recent studies concluded that based on these facts HN may have been considerably (∼40%) less fit compared to HS (Harris and Nielsen, 2016). To capture this asymmetry, I include a 50% lower carrying capacity (K_n_=0.5 K_s_) for Neanderthals. This means that, in equilibrium the same food resources per grid box would have supported a HN population only half of the size of an equilibrium HS population. This asymmetry is also supported by analyses that suggests that the potential habitat range of HN may have been considerable smaller and more fragmented than for HS (Melchionna et al., 2018). Additional experiments with different competition factors β_NS,_ β_SN_ will further elucidate the asymmetry effects between HN and HS. The scaling K_s_(x,y,t)=0.12 N(x,y,t) /(kgC m**^−^**^2^ yr**^−^**^1^ km^2^) is estimated from a constraint that requires the simulated global population of HS in BSL to lie in the range of **2-4 million** individuals at the onset of the Holocene (Figure 2e). Previous studies (Currat and Excoffier, 2011) conducted sensitivity experiments with an even lower K_n_/K_s_ ratio of 25%. Using K_n_=0.5 K_s_ here and for hypothetical constant climate parameters, competitive exclusion (Gilpin et al., 2016; Neuhauser and Pacala, 1999; Roberts and Bricher, 2018; Steele, 2009) and therefore extinction of Neanderthals would eventually occur when *β*_*NS*_ > 0.5. However, in the HDM carrying capacity and growth rate are changing continuously. It is therefore not straight-forward to apply the idealized competitive exclusion criterion (Steele, 2009) to Fisher-Kolmogorov reaction diffusion systems with time-varying parameters.

**Figure. 2.**
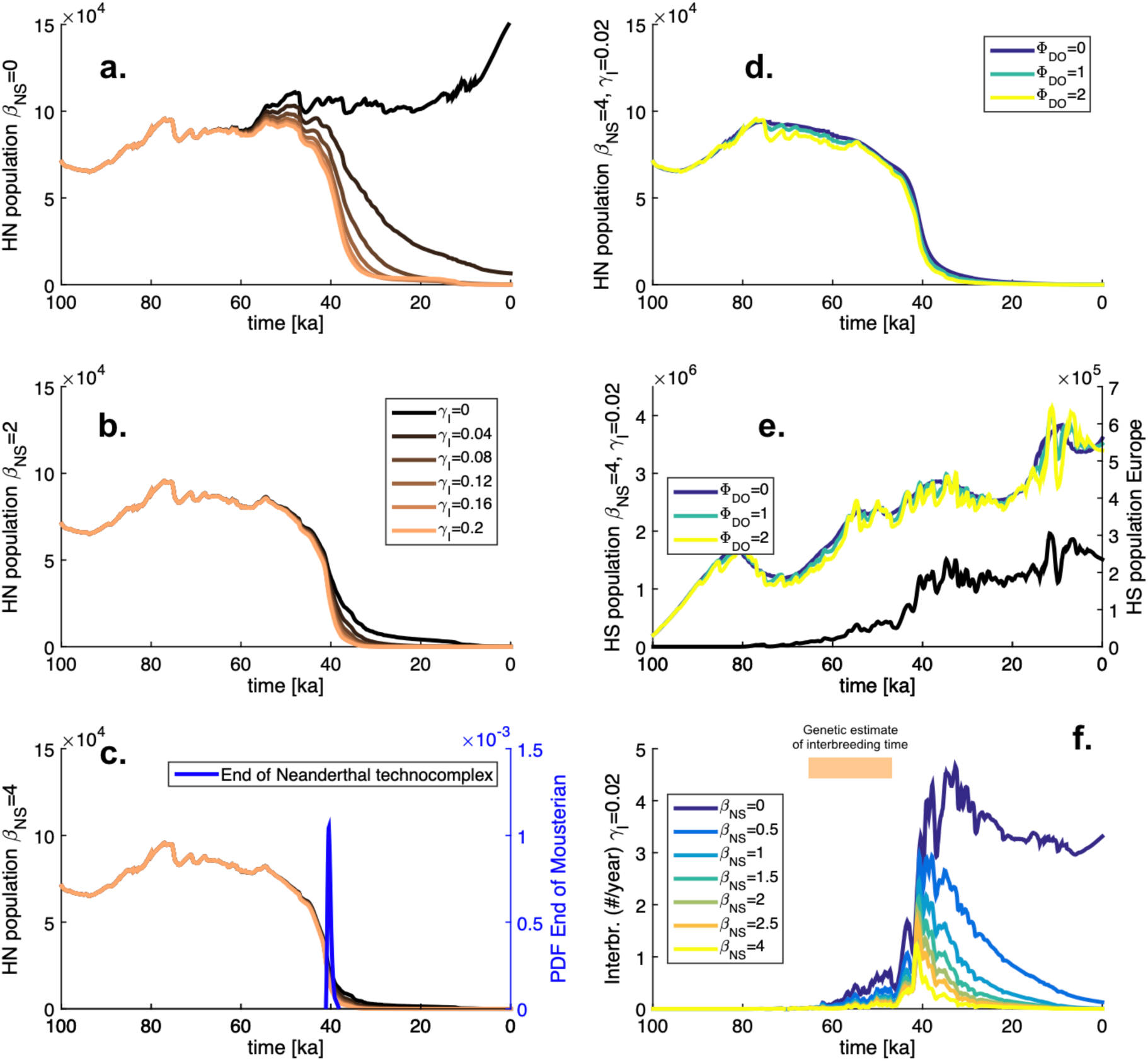
Neanderthal extinction and its parameter sensitivity: a. Global population of HN for different values of interbreeding factor (γ_I_), excluding competitive effects of HS on HN (β_NS_=0). The simulation for γ_I_ =0 and β_NS_=0 (dark brown) is a highly idealized simulation that shows that, in the absence of hominin interactions, climatic effects alone would not have been sufficient to cause a HN extinction; b. Same a., but for moderate competition levels β_NS_=1.5; c. same as a; but for strong competition β_NS_=4. Blue curve represents radiocarbon-based end of Mousterian techno-complex(Higham et al., 2014); d. Global population of HN using different amplitudes of Dansgaard Oeschger variability (Φ_DO_=0,1,2) and realistic interbreeding rates (γ_I_=0.02) and strong competition β_NS_=4 (DOS); e. Same as d., but for global population of HS; black solid line corresponds to ensemble mean HS population size in Europe (STD simulation); f. globally integrated interbreeding rate (individuals per year) for different values of competition and for realistic interbreeding factor γ_I_=0.02 (ENS simulation).

The diffusion terms for Neanderthals and *Homo sapiens* are parameterized as v_n_(x,y,t)=[v_n0_+v_c_ R(x,y) +v_c_C(x,y,t)]ζ(x,y) and v_s_(x,y,t)=[v_s0_+v_c_ R(x,y) +v_c_C(x,y,t)]ζ(x,y) where v_n0_=17, v_s0_=27 km^2^/year represent the background diffusivities for HN and HS (Table 1). The diffusion rates chosen here are in the broad range suggested by Young and Bettinger (1992, 1995). The second and third terms capture the effect of increased diffusion along rivers (captured by stationary river mask R(x,y) and time-evolving coastlines [C(x,y,t)]. ζ(x,y) is a fixed surface roughness mask, which assigns values of 1 to low topographic roughness areas and values close to zero to high topographic roughness regions. v_n_(x,y,t), v_s_(x,y,t) for t=100 ka are depicted in Figure 3a,b.

**Figure. 3.**
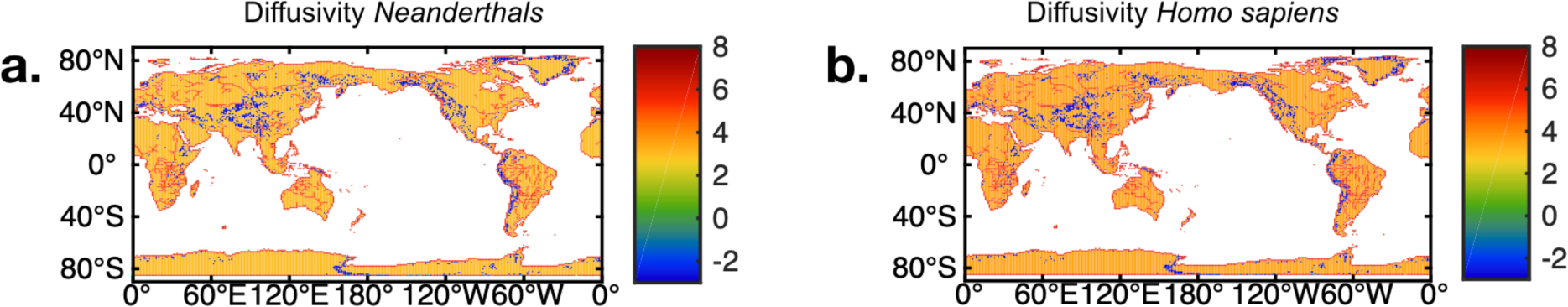
Spatial pattern of diffusivity: Logarithm of diffusivity (km^2^/year) at 100 ka of Neanderthals (a) and Homo sapiens (b), representing a different background value and the possibility for enhanced coastal and river migration speeds. A slowdown effect in regions of rough topography is also included.

**Table 1.** Ensemble model experiments conducted, and key parameter values used. Throughout the simulation, the coastline C(x,y,t) (land-sea distribution) varies temporally, following estimates of past sea-level variability (Waelbroeck et al., 2002) (**Supplementary movies 1-4**).

| Model simulation | Start time [ka] | $\beta_{NS}$ | $\beta_{SN}$ | $\gamma_i$ | $\Phi$ | $V_{n0}$ [1/km<sup>2</sup>] | $V_{s0}$ [1/km<sup>2</sup>] | $\alpha_{n0}$ [%/year] | $\alpha_{s0}$ [%/year] | $T_{on}$ [C] | $T_{os}$ [C] | $\sigma_n$ [C] | $\sigma_{so}$ [C] |
| --- | --- | --- | --- | --- | --- | --- | --- | --- | --- | --- | --- | --- | --- |
| Baseline<br>Simulation, no<br>competition, no<br>interbreeding<br>(BSL) | 100 | 0 | 1 | 0 | 2 | 17 | 27 | 0.021 | 0.12 | 13 | 29 | 8 | 150 |
| Interbreeding<br>Simulations, no<br>competition<br>(IBR), | 100 | 0 | 1 | 0-0.2 | 2 | 17 | 27 | 0.021 | 0.12 | 13 | 29 | 8 | 150 |
| Ensemble<br>Simulations,<br>combined<br>interbreeding and<br>competition<br>(ENS), | 100 | 0-4 | 1 | 0-0.2 | 2 | 17 | 27 | 0.021 | 0.12 | 13 | 29 | 8 | 150 |
| Standard<br>Simulation,<br>Realistic<br>interbreeding and<br>competition<br>(STD) | 100 | 4 | 1 | 0.02 | 2 | 17 | 27 | 0.021 | 0.12 | 13 | 29 | 8 | 150 |
| Dansgaard<br>Oeschger<br>simulations,<br>realistic<br>interbreeding and<br>competition<br>(DOS), | 100 | 4 | 1 | 0.02 | 0,1,<br>2 | 17 | 27 | 0.021 | 0.12 | 13 | 29 | 8 | 150 |

In addition to the carrying capacity-climate dependence, the coupling of demographic processes with the time-varying climate environment is captured by a temperature-dependence of the growth rates, expressed as α_n_(x,y,t)= α_n0_ e^[(T(x,y,t)-Ton)^2/σn^2]^ and α_s_(x,y,t)= α_s0_ e^[(T(x,y,t)-Tos)^2/σs(t)^2]^ ε(t) where T(x,y,t) represents the simulated annual mean LOVECLIM surface temperature field in °C and α_n0_, α_s0_ are the maximum annual growth rates, which I choose as 0.021% and 0.12%/year, respectively. An asymmetry in these parameters (Table 1) is further supported by the fact that the genetic diversity, as well as the effective population of Neanderthals was considerably lower than for HS (Castellano et al., 2014). It allows HS to expand faster into territories than HN, in accordance with the larger potential HS habitat argument (Melchionna et al., 2018). Such processes may become relevant for the reoccupation of grid boxes during a Dansgaard-Oeschger stadial/interstadial transition. T_on_=13°C and T_os_=29°C represent optimal annual mean habitat temperatures for HN and HS, respectively and σ_N_=8. *ε*(t) is 1 at the beginning of the simulation and increases linearly to 1.5 towards the end of the simulation to capture an overall acceleration of dispersal and a general adaptation as well as social and cultural evolution of HS. Furthermore, I allow σ_s_ to vary in time to capture the fact that the habitat range of HS broadened as a result of mutations (Beleza et al., 2013) that occurred between 42-38 ka which allowed Eurasian HS to down-regulate melanin production, develop lighter skin and synthesize Vitamin D in higher latitudes with less solar radiation. This is parameterized through σ_s_ (t) = σ_so_ atan [0.2 (t_m_ -t)]/π + 80, where time t is measured in ka and I choose σ_so_=150 and t_m_=40 ka. Simulations, that do not include this process (not shown) exhibit unrealistic (too early) arrival times of HS in Europe and corresponding early Neanderthal extinction. I have therefore decided not to include these experiments in the overall parameter sensitivity studies described here. To capture the Allee effect, which describes the fact that small groups are more vulnerable and more likely to go extinct (Dennis, 1989), I assume that HN and HS were not able to survive in areas with a desert fraction > 90%, or if their population density drops below a critical density per grid cell (<10^-6^ individuals/km^2^). This criterion becomes active in the ice-sheet-covered regions in my simulation (Ganopolski et al., 2010), low elevation deserts, such as parts of the Sahara during Northern Hemisphere summer aphelion conditions and high-altitude deserts such as the Tibetan Plateau. The time-varying desert-fraction is as one of the climatic boundary conditions described in the next paragraph. Important parameters of the HDM2 are listed in Table 1 for the individual experiments.

To represent existing knowledge on habitats, genetic diversity, effective population sizes, I included several demographic asymmetries between HN and HS into the HDM. This concerns parameters for initial population size, diffusivity, growth rates and carrying capacity. However, these asymmetries alone do not necessarily lead to a realistic HN extinction in my simulations. Therefore, additional sensitivity experiments are conducted by varying competition and interbreeding parameters as well as the effect of DO events. This allows me to quantitatively test previously suggested hypotheses on HN extinction in a more realistic setting.

Equations (1) and (2) are discretized with an explicit finite difference scheme for diffusion equations with inhomogeneous diffusivity (Praprotnik et al., 2004) and by using a timestep of 10 years. One 100,000-year-long simulation takes about 2 minutes on a single Intel Skylake node with 768 GB memory and 2TB NVMe SSD.

### 2.2 Climate forcings and boundary conditions

#### 2.2.1 Orbital-scale climate model simulation

The effects of slowly evolving orbital-scale/glacial boundary conditions on the climate system [T_orb_(x,y,t) N_orb_ (x,y,t), D_orb_ (x,y,t)], are included into the HDM2 by implementing the simulated fields from a transient glacial-interglacial climate model simulation conducted with the earth system model LOVECLIM (abbreviated as ORB for orbital-scale) (Timmermann and Friedrich, 2016). The simulation ORB is based on the earth system model LOVECLIM, version 1.1 which uses the three-level T21 atmospheric model ECBilt. The model contains a full hydrological cycle, which is closed over land by a bucket model for soil moisture and a runoff scheme. Diabatic heating due to radiative fluxes, the release of latent heat and the exchange of sensible heat with the surface are parameterized. The ocean-sea ice component CLIO consists of a free-surface Ocean General Circulation Model with 3°x3° resolution coupled to a thermodynamic-dynamic sea ice model. Coupling between atmosphere and ocean is established through the exchange of freshwater and heat fluxes. The terrestrial vegetation module VECODE, computes the annual mean evolution of the vegetation cover (tree, desert and grass fractions) based on annual mean values of several climatic variables, including precipitation and temperature.

Orbital-scale time-evolving ice-sheet boundary conditions in ORB are prescribed by changing ice-sheet orography and surface albedo. The corresponding anomalies were derived from the time-dependent ice-sheet reconstruction obtained CLIMBER earth system model of intermediate complexity (Ganopolski et al., 2010). In LOVECLIM also the vegetation mask is adjusted to reflect time-evolving changes in ice-sheet covered areas. Time-varying atmospheric greenhouse gas concentrations are obtained from the EPICA DOME C ice core (Luthi et al., 2008). I also adopt orbitally-induced insolation variations (Berger, 1978). The forcing is applied to LOVECLIM using an acceleration technique, which compresses the time-varying external boundary conditions by a factor of 5. Instead of running the coupled model for the entire period of 784,000 years the model experiment is 156,800 years long, while covering the entire forcing history of the last 784 kyr. Our current model version has a higher climate sensitivity (Timmermann and Friedrich, 2016), which amounts to ∼4°C/CO_2_ doubling. The result is a more realistic glacial/interglacial amplitude in surface temperatures compared to palaeo-proxy data. The ORB model simulation does not include the effects of millennial-scale variability associated with Dansgaard-Oeschger and Heinrich events. This variability is included through a secondary model/data-based procedure (Figure 1).

#### 2.2.2 Millennial-scale variability

Whilst the full climate model simulation covers 784 ka, only the last **100** ka are used in this study to force the HDM2 (see Figure 2). The variables that will be used as part of the climate forcing of the HDM2 are the simulated changes in temperature T_orb_ (x,y,t), which is needed for the calculation of growth rates α_n_, α_s_, the net primary productivity N_orb_ (x,y,t) which is used to calculate the carrying capacities for HN and HS [K_n_(x,y,t), K_s_(x,y,t)] and the desert fraction D_orb_ (x,y,t), which gets translated into instant mortality for HN and HS if the value exceeds 90%. To additionally include the effects of millennial-scale Dansgaard-Oeschger (DO) variability into the HDM2, I reconstructed the corresponding anomalies of surface temperature T_m_(x,y,t), and net primary productivity N_m_(x,y,t) and desert fraction d_m_(x,y,t), where the subscript _m_ stands for the millennial-scale climate anomalies obtained from a previously conducted transient LOVECLIM climate model simulation (Menviel et al., 2014). This simulation is a climate model-based hindcast covering the period from 50-30 ka, which includes both orbital scale forcings (ice-sheets, greenhouse gas concentrations and orbital changes) and freshwater-forced millennial-scale climate shifts associated with each observed DO and Heinrich event during this period. For this climate model simulation I calculated the regression patterns P_N_(x,y), P_T_(x,y), P_D_(x,y) between a simulated temperature index on the Iberian Margin T_I_(t) and the corresponding spatio-temporal fields:

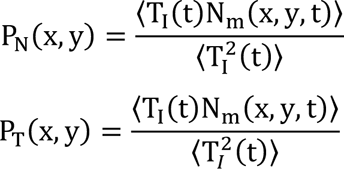

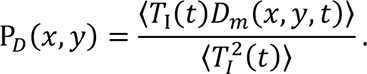

To reconstruct the millennial-scale anomalies [T*_m_’(x,y,t), N*_m_’(x,y,t), D*_m_’(x,y,t)] for the entire HDM2 period 100-0 ka, I multiply the model-based regression patterns with a high-pass filtered normalized temperature index, which characterizes Dansgaard Oeschger events. In this case it is a high-pass filtered version of a synthetic Greenland temperature record (Barker et al., 2011) T_G_’, which captures DO variability extending back to 784 ka. One therefore obtain:s T*_m_’(x,y,t)= T_G_’(t) P_T_(x,y); N*_m_’(x,y,t)= T_G_’(t) P_N_(x,y); D*_m_’(x,y,t)= T_G_’(t) P_D_(x,y). The resulting millennial-scale anomaly maps are then added back to the corresponding fields of the orbital scale SIM simulation [T_ORB_(x,y,t), N_ORB_(x,y,t), N_ORB_(x,y,t)] to give T(x,y,t)= T_ORB_(x,y,t) + Φ T_M_(x,y,t); N(x,y,t)= (N_ORB_(x,y,t) +Φ N_M_(x,y,t)) Θ[N_ORB_(x,y,t) +Φ N_M_(x,y,t)] (Heaviside function prevents terms from becoming negative); D(x,y,t)= [D_ORB_(x,y,t) + Φ N_D_(x,y,t)] Θ[D_ORB_(x,y,t) + Φ N_D_(x,y,t)] Θ[100-(D_ORB_(x,y,t) + Φ N_D_(x,y,t))]) (product of Heaviside functions prevents terms from becoming negative or exceeding 100%).

Our approach generates climatic timeseries at every grid point that show close resemblances with various paleo-climate reconstructions from around the world (Fig. 4).

**Figure 4.**
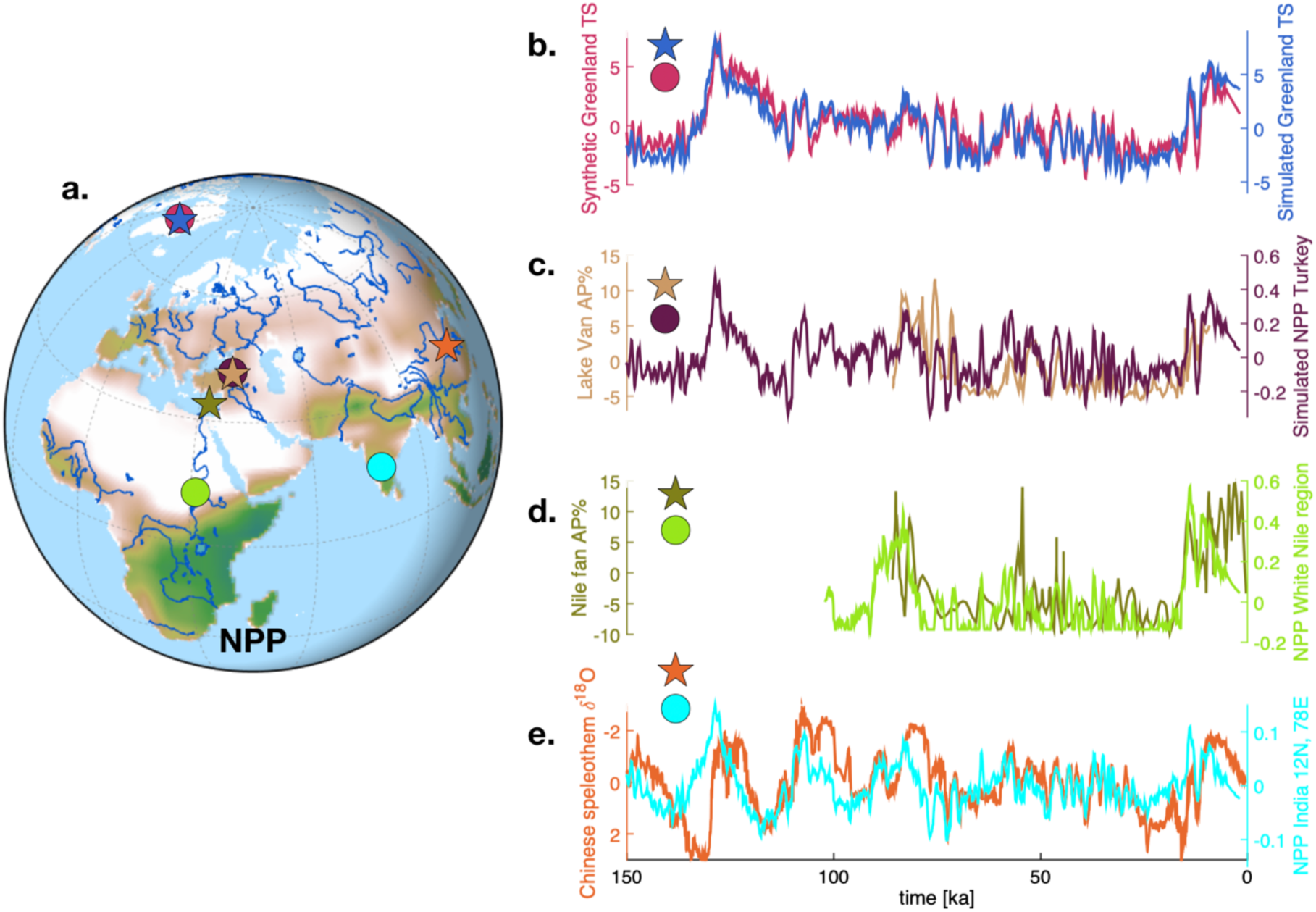
Comparison between climate forcing and paleo-proxy data: a. location of timeseries extracted from model (circles) and for proxy data location (stars); b. Temperature anomalies in Greenland from climate forcing reconstruction and synthetic Greenland proxy record(Barker et al., 2011); c. simulated anomalies of net primary production and arboreal pollen (Litt et al. 2014) from Lake Van; d. Simulated anomalies of net primary production in White Nile area and arboreal pollen extracted from sediment core MD9505 from the Nile fan (Langgut et al., 2018); f. simulated anomalies of net primary production in Southern India, representing changes in Indian summer monsoon and δ^18^O composite record from Sanbao and Hulu caves in China (Wang et al., 2008b), which may capture a downstream effect of the Indian Summer monsoon.

The comparison between reconstructed and observed Greenland temperature variability for the standard runs with Φ=2 shows an excellent agreement, on both orbital and millennial timescales (Figure 4b). Note, that the timing of the Greenland high-frequency millennial-scale variability (Barker et al., 2011) is included as a forcing for the hybrid climate reconstruction, thus explaining the high correlation on millennial timescales. However, the orbital-scale contribution of the HDM climate forcing originates directly from the LOVECLIM transient 784 ka model simulation. It is therefore not constrained through empirical proxy data shown in Figure 4. Figure 4 also shows a good agreement on both timescales between reconstructed and paleo vegetation variations in the Levant region (Litt et al. 2014) (Figure 4c). To further highlight the fidelity of the climate forcing in areas relevant for early hominin dispersal, I compare arboreal pollen data carried downstream by the Nile river (Langgut et al., 2018) with the simulated vegetation variations farther upstream in the White Nile region (Figure 4d). A similar non-local comparison is made for Chinese speleothem δ^18^O variations (Wang et al., 2008a) and simulated rainfall variations in India (Figure 4e). For this comparison it is assumed that the δ^18^O of speleothem calcite does not represent local rainfall in China, but rather a remote signal which switches on precessional timescales between the Indian summer Monsoon and local Pacific moisture sources (Maher and Thompson, 2012). Other geographic and climatic features, that may have changed hominin dispersal routes temporarily, such as pro-glacial lakes, volcanic eruptions, rapid flooding of coastlines during Heinrich events are not included in the present form of the climate forcing.

### 2.3. Experiments

The dispersal model is configured to ascertain the effects of resource competition, interbreeding, demographic factors and abrupt climate change on the demise of the Eurasian Neanderthal population. An ensemble of parameter sensitivity experiments was conducted to capture these processes (Table 1). Within this ensemble I look for reasonable parameter configurations that yield realistic simulations of HN, HS demography and which are qualitatively consistent with archeological, anthropological, demographic and genetic data.

An idealized baseline simulation (**BSL**) of the HDM is conducted with realistic climate and vegetation forcing, time-evolving coastlines, a gradual increase of HS growth rate to represent adaptation to a variety of environmental conditions and in particular to reduced sunlight in higher latitudes (Beleza et al., 2013; Crawford et al., 2017). BSL ignores the effect of HS on HN (competition and interbreeding parameters are set to β_NS_=0, γ_i_=0, respectively) (**Supplementary movie 3**). I also conduct a series of parameter perturbation simulations to explore the sensitivity of the HDM solutions to a variety of interbreeding rates ranging from 0 to unrealistically large values of 20% (**IBR**).

Even though competition is represented in the HDM2 as resource competition (Aoki et al., 1996; Okubo et al., 1989; Young and Bettinger, 1992), it captures a wider range of competitive advantages that AMHs may have had over Neanderthals, such as e.g. in cognitive and social abilities (Kochiyama et al., 2018), cultural superiority (Gilpin et al., 2016) or higher resistance to pathogens from Africa (Houldcroft and Underdown, 2016). In the HDM2 their combined effect is included in one parameter (β_NS_), which quantifies the ratio of AMH “superiority” over Neanderthals, expressed in terms of resource competition. Furthermore, I have assumed a higher AMH fecundity as compared to Neanderthals (Roberts and Bricher, 2018) (Table 1).

To further test the sensitivity of HDM2 to the competition and interbreeding parameters, another ensemble of experiments (**ENS**) was conducted, in which both parameters key parameters are changed simultaneously: β_NS_=0-4, γ_i_=0-0.2. This ensemble uses realistic climate forcing (Φ=2). Among the ensemble members one simulation is picked which includes realistic values of competition, as well as interbreeding (β_NS_=4, γ_I_=0.02) and agree well with demographic, genetic and archaeological datasets. This simulation will be referred to as Standard Run (**STD**). To test the effect of Dansgaard-Oeschger variability on HS, HN I run simulations with Φ=0,1,2 and for realistic values of competition and interbreeding. These experiments will be referred to as **DOS**.

In the simulations presented here, I tried to constrain initially as many parameters as possible from empirical evidence. These include demographic parameters K_n_, K_s_, v_n_, v_s_, α_n_, α_n_, the time of skin color mutation t_m_, as well as potential habitat ranges, characterized by T_on_, T_os_, σ_n_, and σ_s_ and initial conditions for HN (Bocquet-Appel and Degioanni, 2013) and HS (Sjodin et al., 2012). The most uncertain parameters (competition β and interbreeding levels γ_i_) are then further perturbed to determine key sensitivities or test specific hypotheses on HN extinction (DO events, Φ) (see Table 1). It should be noted here, that already in the initial parameter choices I made and in agreement with a plethora of previous studies, an asymmetry between HN and HS is present (e.g. K_n_ <K_s_), which would lead to a long-term demise of the weaker (HN) population. The key goal of my sensitivity experiments is to determine under which demographic and climatic conditions, the HDM simulates a realistic HN extinction in terms of both, time and space.

## 3. Results

### 3.1 Resource competition

Starting at 100 ka with initial populations of 200,000 HS (Sjodin et al., 2012) in eastern Africa and 70,000 HN (Bocquet-Appel and Degioanni, 2013) in Eurasia, the Baseline experiment (**BSL**) simulates an overall growth of both HN and HS populations until **50 ka**, a relatively stable plateau during parts of Marine Isotope Stage 3 (MIS3, 57-29 ka,) and a second phase in population growth during the last glacial termination and into the early Holocene from ∼20 ka −8 ka (Fig. 2a, e), during which HS reaches global populations levels of about 4 million individuals. Demographic data simulated during the Holocene will not be interpreted here, because the model does not properly capture non-hunter and gatherer populations. The choice of lower initial population sizes is inspired by recent genetic studies that suggest a considerably reduced effective population size of HN relative to HS (Castellano et al., 2014). From 60-40 ka the typical density of HN (HS) in Europe simulated by this idealized experiment ranges geographically from 0.01-0.025 (0.04-0.07) individuals/km^2^ (Figs. 2, 5), consistent with other studies (Currat and Excoffier, 2011).

**Figure. 5.**
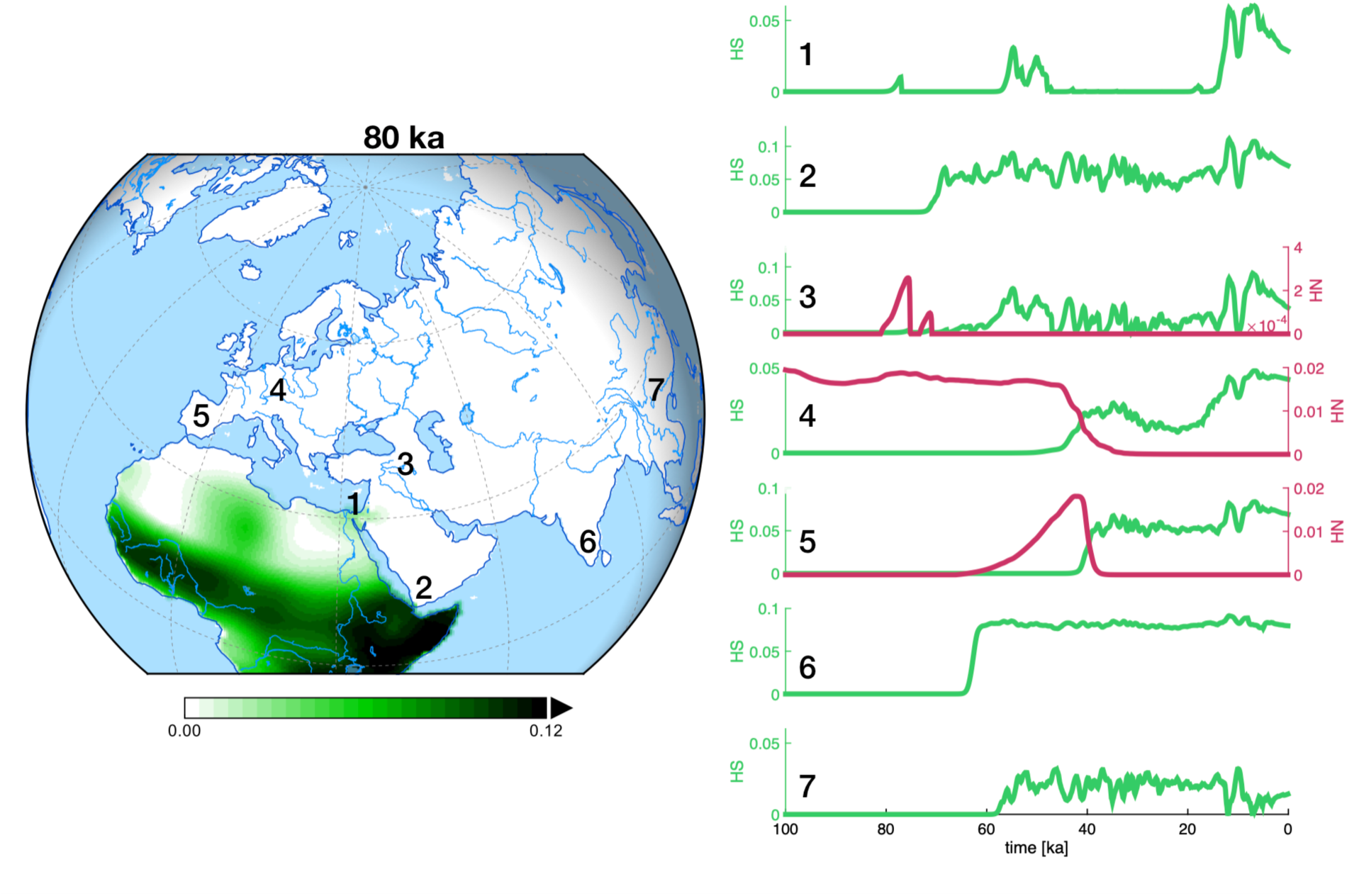
Regional Population evolution: Regional changes in population density (individuals/km^2^) of AMH (HS) (green lines) and Neanderthals (HN) (red lines) at key sites, indicated by numbers on map (left). Shading in left panel shows HS population density at 80 ka for a 100 ka initial condition (sea level conditions have not been adjusted on map). Note, that a 125 ka initial condition would yield an earlier migration wave out of Africa, which leads to a 100 ka dispersal wave from Africa into Eurasia and more realistic, earlier, arrival times in Eastern Asia (Timmermann and Friedrich, 2016) around 80-60 ka.

During MIS3, the simulated *Homo sapiens* population size in Europe amounts to about 200,000 individuals (Figure 2e), which also agrees well with recent model-based estimates (Tallavaara et al., 2015). We also see that Dansgaard-Oeschger variability can influence the size of the European HS population by about +/-15%. The second growth phase of HS population size from 20-8 ka is mostly related to a rapid expansion of HS into high latitudes and into North America. Annual growth rates during major phases of population expanse and during Dansgaard-Oeschger interstadials attain values of about 0.02-0.04%, which agrees with previous empirical estimates for the Holocene (Zahid et al., 2016) of 0.03-0.06%. The HDM2 experiments simulate a first HS dispersal from Africa into Eurasia through the Sinai Peninsula around 80 ka, followed by a second wave from 59-47 ka (**Supplementary movie 2,** Fig. 5**, left panel**).

In a series of experiments, the sensitivity of global Neanderthal extinction dates is calculated with respect to the competition factor β_NS_. The BSL scenario without competition shows no extinction of HN, in spite of the presence of Dansgaard-Oeschger variability in this scenario (Figure 2a**, black line**). By introducing a reasonable level of food resource competition (β_NS_=1.5; β_SN_=1) in the carrying capacity term (**ENS**) and ignoring interbreeding (γ_i_=0), HN extinction, defined here as the time when the global HN population size drops below 3000 individuals, occurs around 10 ka (Figure 2, 6b).

**Figure 6.**
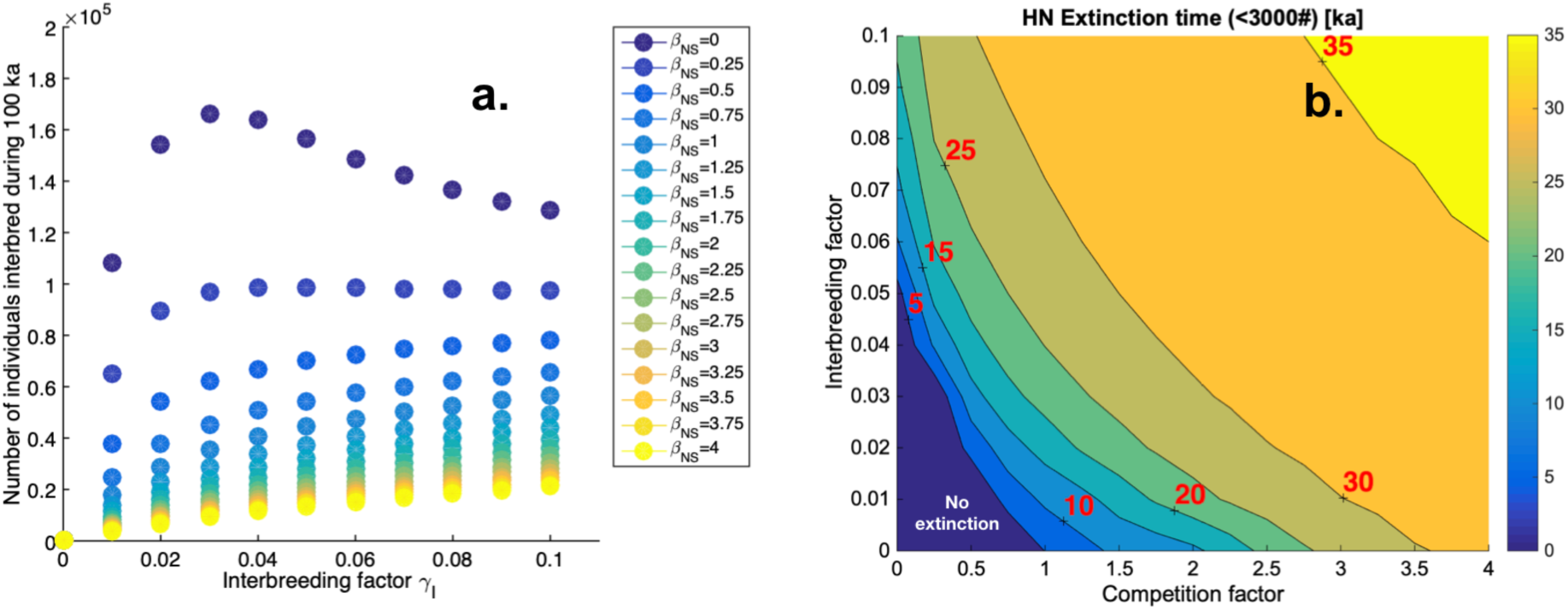
Demographic effects on Neanderthal disappearance: a. total number of interbreeding individuals integrated for entire simulation and globally for different values of competition ad interbreeding; b. Extinction time of HN in ka (defined as the time when the global population of HN drops below 3000 individuals) for different values of competition and interbreeding factor. For such very population numbers strong Allee effects could easily drive populations into extinction, even earlier.

A more aggressive choice, in which HS is assumed to have considerable advantages over HN (as expressed by β_NS_=3-4; β_SN_=1), shows a much earlier HN extinction (Figs. 2, 6**, Supplementary movie 1**) around 25-32 ka, which clearly illustrates the power of competitive exclusion in multi-population systems. Based on the model equations (1,2), one can immediately see that by ignoring the inhomogeneous diffusion and interbreeding terms, the coupled Fisher Kolmogorov model (Methods) reduces to a coupled Lotka-Volterra competition system, in which HS in the equilibrium solution eventually replaces HN when β_NS_>K_N_/K_S_=0.5 and β_SN_=1< K_S_/K_N_=2 (Neuhauser and Pacala, 1999; Steele, 2009). Calculations of HN extinction times calculated with the full model as part of **ENS** and for different values of β_NS_ (Figure 7) are qualitatively consistent with this mathematical extinction threshold, even though the temporal forcing (non-autonomous system), and the 2-dimensional diffusion terms complicate the overall dynamics in HDM2, relative to the zero-dimensional model. Another ensemble for varying competition factors was run for Φ=0 (not shown), showing qualitatively similar results in terms of extinction times as in Figure 7, but marked differences in the millennial-scale fluctuations of HN global population size. It should be noted here that this study goes beyond theoretical considerations on the asymptotic extinction limit of a less competitive agent. The experiments presented here address a much more specific and difficult question: which parameter choices (corresponding to different HN extinction hypotheses) lead to a realistic extinction in the HDM in terms of timing and spatial extent.

**Figure 7.**
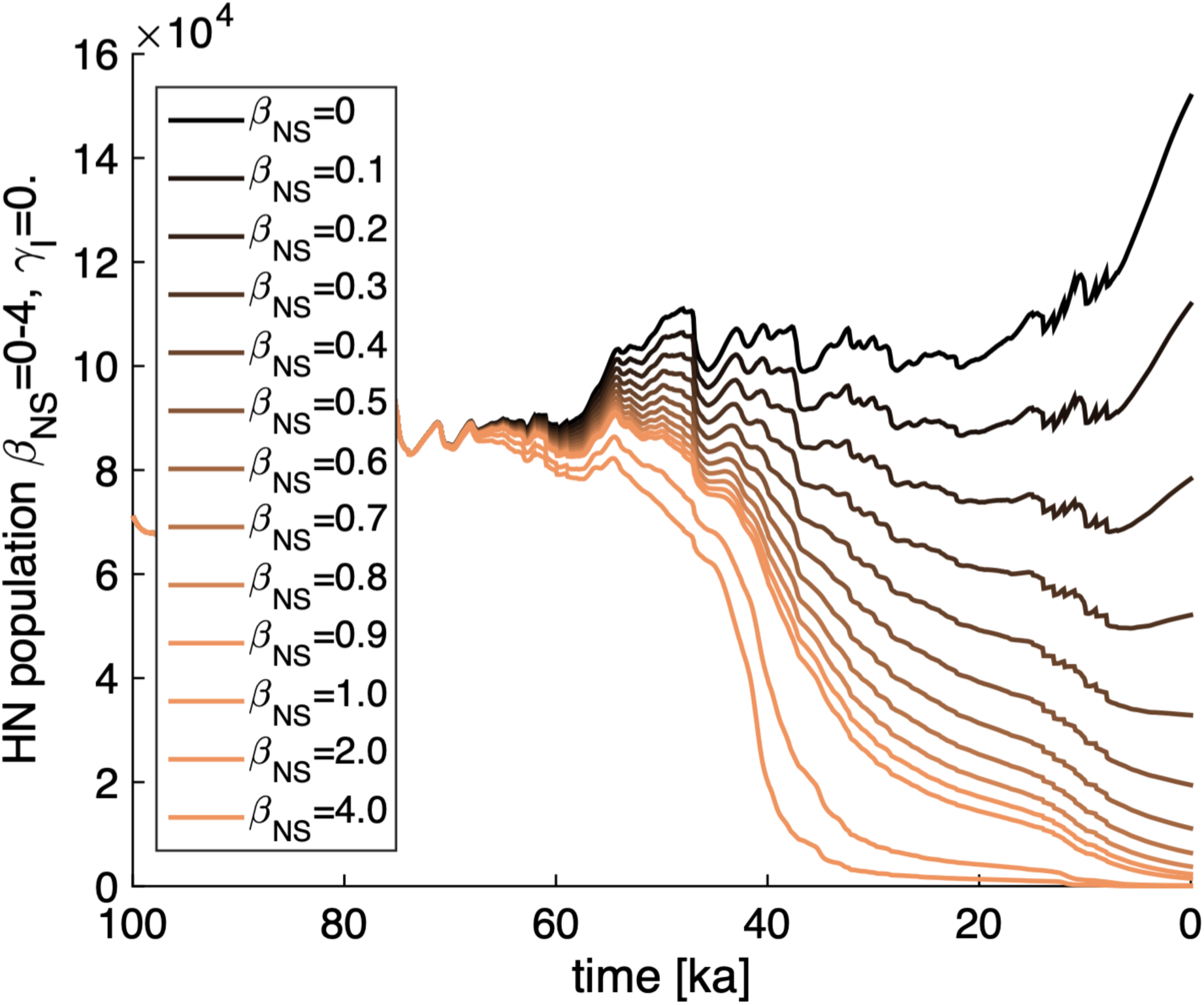
Extinction time dependence on competition: Simulated HN population size for different values of competition β_NS_=0-4, but no interbreeding (subset of **ENS** simulation). The theoretical threshold for extinction in a zero-dimensional coupled Lotka-Volterra model version without interbreeding, inhomogeneous diffusion and for constant carrying capacity and growth rate is β_NS_=0.5. The dark brown line corresponds to a simulation, in which HN can develop without interacting with HS.

### 3.2 Interbreeding

Interbreeding may have been one other crucial factor in the interaction between HS and HN during MIS3. First evidence for interbreeding and hybridization can be traced back to at least 47-65 ka (Fu et al., 2015; Fu et al., 2014; Sankararaman et al., 2012) (Fig. 2f). The exact rates of interbreeding still remain highly uncertain. According to recent modeling studies (Currat and Excoffier, 2004; Neves and Serva, 2012) they may have been significantly less than 2%, which corresponds to a few hundred to a few thousand interbreeding individuals in total during the entire time. It should however be noted here, that the recent discovery of a Neanderthal-Denison-F1 hybrid (Slon et al., 2018) supports the notion that interbreeding among hominins may have been much more common than previously thought and may have even occurred multiple times in different areas (Villanea and Schraiber, 2019). The interbreeding simulations (**IBR, ENS**), in which the interbreeding rates were changed from 0 to unrealistically large values of 20% (Fig. 2 a-c), show that interbreeding plays a much bigger role for low resource competition (Fig. 6b). As the effect of competitive exclusion increases, the role of introgression (Fig. 2 f) is reduced and extinction occurs within a time window from 25-35 ka (Fig. 6b). Total numbers of interbreeding individuals during the simulation constrain the combination of the values of the interbreeding rate γ_I_ and the competition factor β_NS_ (Fig. 6a). It is found that the weak genetic introgression constraint for the total number of successfully interbreeding individuals (<10,000) can only be met for γ_I_<0.02 and β_NS_>2. These values translate into likely HN extinction times of >20 ka (Fig. 6b). A more detailed view on the dynamics of interbreeding simulated in the model for more realistic parameter values (β_NS_=4, γ_I_=0.02), referred to as Standard Run (**STD**), shows a more complex pattern of assimilation of HN either into HS groups or vice versa (Figs. 8, 9 **Supplementary Movie 4**).

**Fig. 8.**
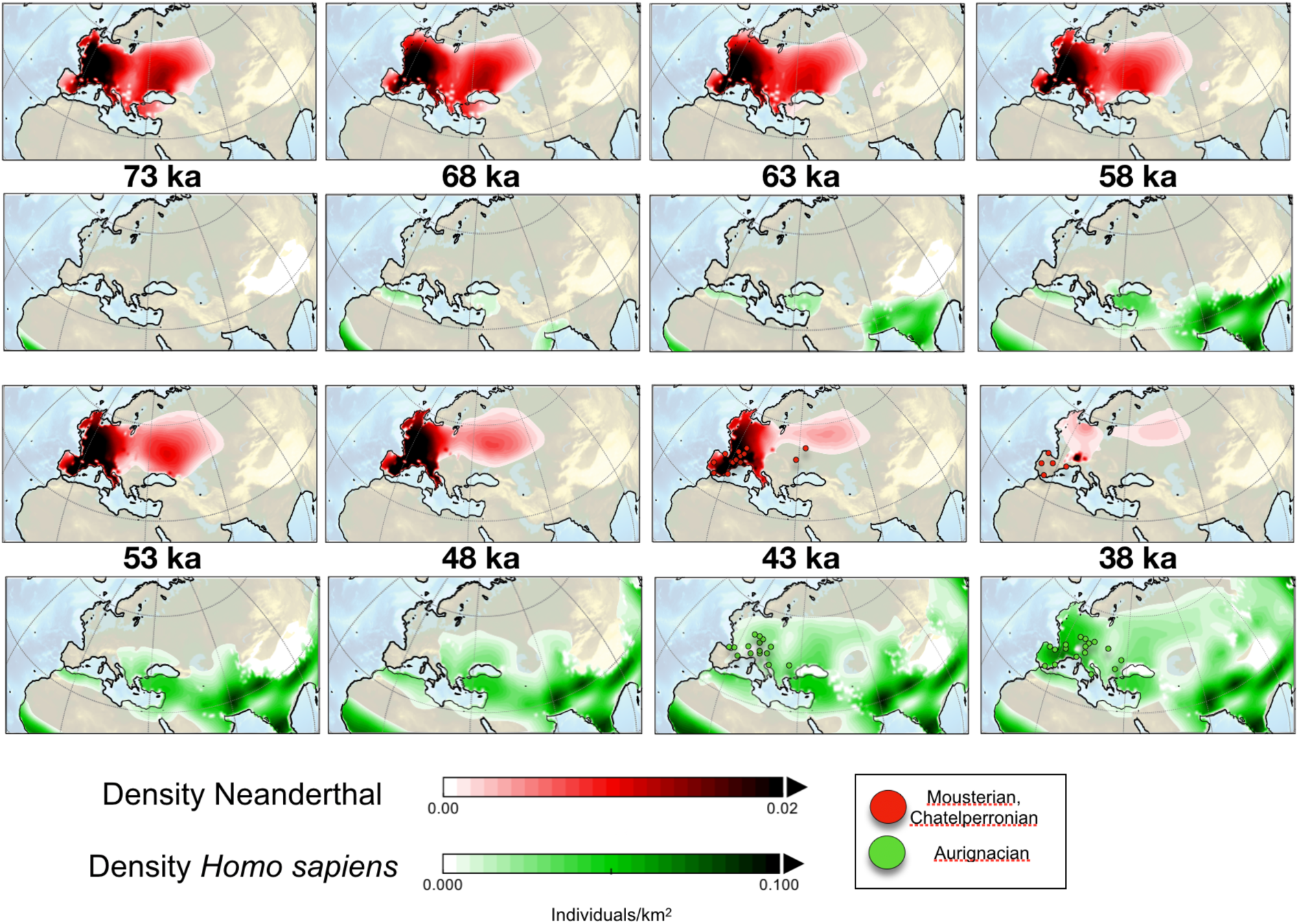
Population growth of Anatomically Modern Humans and Neanderthals for realistic competition and interbreeding scenario: Evolution of HN population density (individuals/km^2^) (red shading), simulated by Standard run (**STD**) for snapshots at 73, 68, 43, 58, 53, 48, 43, 38 ka; Green shading, same as red, but for HS population density; lower right panels: sites of radio-carbon-dated Mousterian, Châtelperronian and Aurignacian techno-complexes are included (Banks et al., 2008) (aggregated over 5 ka windows).

**Fig. 9.**
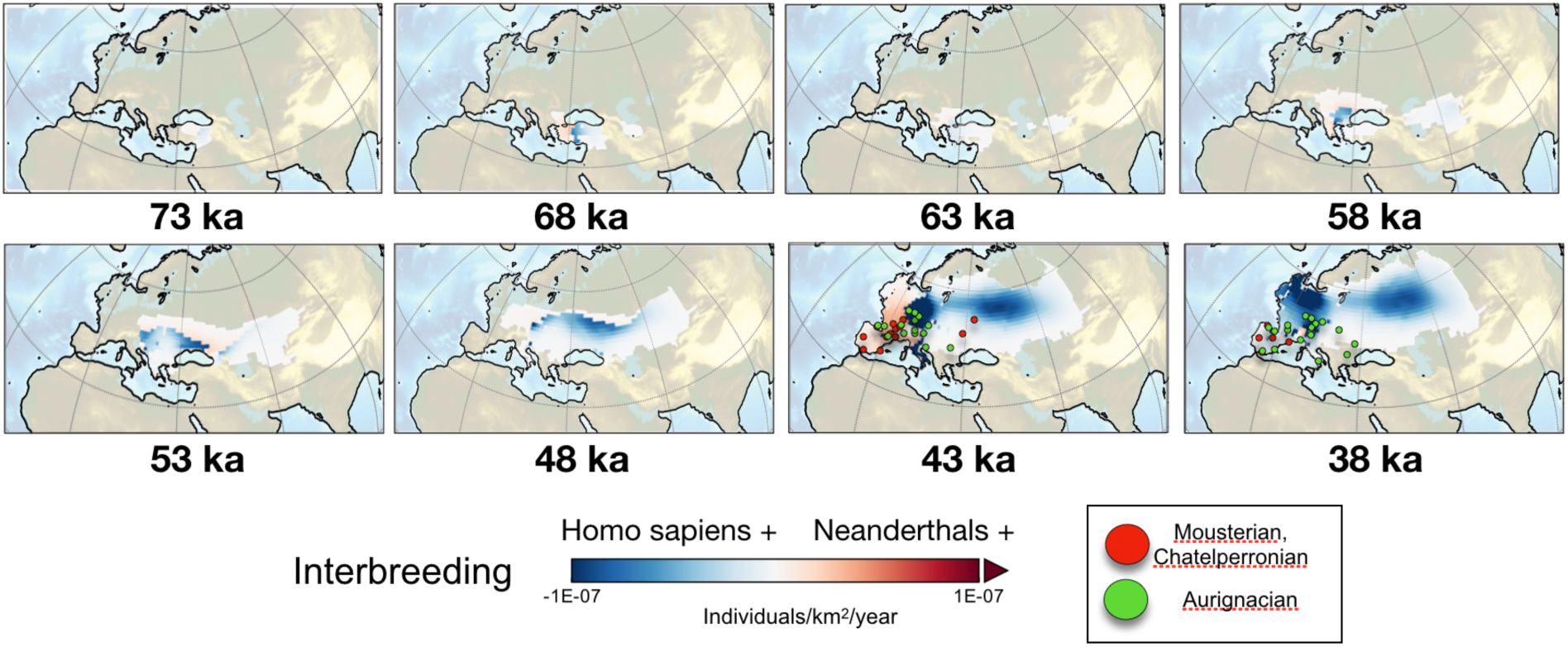
Propagation of Hominin interbreeding front: Snapshots for different time period of interbreeding rates [individuals/km^2^/year] for STD simulation using β_NS_ = 4 and γ_i._ = 0.02. Positive (negative) values refer to the case that joint offspring assimilates into the Neanderthal (Homo sapiens) group. Green and dots refer to sites of radio-carbon-dated Mousterian, Châtelperronian and Aurignacian techno-complexes (Banks et al., 2008), aggregated in 5 ka windows. Note, that the model is a continuous model. Values of 10^-7^ individuals/km^2^/year correspond to ∼1 interbreeding event in a 1°x1° grid box every 1000 years.

In STD the demise of HN begins as the first stable HS population establishes in Eastern Europe around ∼50 ka (Figs. 5, 8, 9). Establishing densities at least comparable to those of the HN population in Eurasia, the westward propagating HS wave then reaches Northern France by 43 ka and the Iberian Peninsula by 38 ka (Fig. 8**, Supplementary movie 4**). These simulated dates are in qualitative agreements with fossil data (Fu et al., 2015) (37-42 ka), and the radio-carbon dated Aurignacian sites (Banks et al., 2008) (Figs. 8, 9).

Around 43 ka, the simulated density of HN in Central Europe decreases rapidly, as Neanderthals and AMH begin to compete for the same food resources (Figs. 2, 5) and start to interbreed, which is illustrated by areas of opposing shaded colors (red ρ_N_> ρ_S_; blue ρ_S_> ρ_N,_ see equations (1,2)) (Fig. 9). This is consistent with archeological evidence that suggests that during the period from 43-38 ka, both Mousterian and Châtelperronian (attributed to Neanderthal groups) and Aurignacian (associated typically with *Homo sapiens*) techno-complexes can be found. In the simulation the last HN refugia can be found in the Alps, northern Europe and east of Ural (Fig. 10a). This in contrast with some archeological and anthropological evidence, which places the last refugia on the Iberian Peninsula (Banks et al., 2008) (Fig. 8). However, other studies have argued that the presence of lithic Mousterian technology in the Urals dating back to about 31-34 ka may be an indication for a last refugium of Neanderthals outside southern or central Europe (Slimak et al., 2011), lending further qualitative support for the **STD** simulation (Fig. 8).

**Fig. 10.**
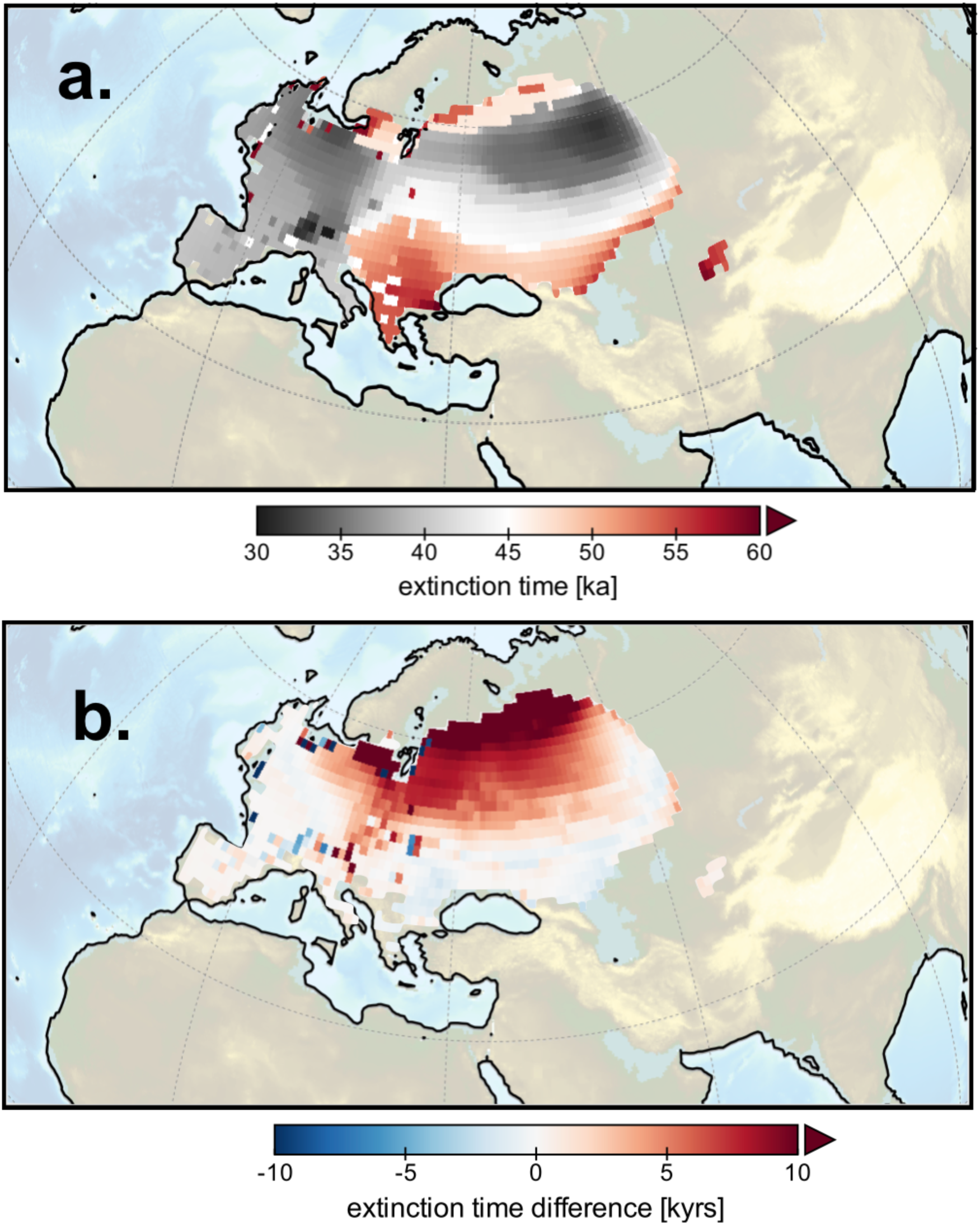
Effect of Dansgaard-Oeschger events on Neanderthal extinction: a. time [ka] when Neanderthal density drops below a minimum threshold value of 0.0001 individuals/km^2^ in STD simulation; b. difference between STD simulation and simulation with same demographic parameters, but without Dansgaard-Oeschger events Φ_DO_=0. Positive values correspond to earlier HN extinction due to DO variability.

It is important to note that there is an initial state-dependence of the simulated arrival times in Europe and a moderate sensitivity with respect to the dispersal velocity of HS *u*_*s*_, which can be approximated as 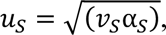, where *v*_*s*_ and *α*_*s*_, represent the average diffusivity and the growth rate of HS, respectively (Table 1).

### 3.3 The role of Dansgaard-Oeschger events

The idea that abrupt climate change, associated with Dansgaard-Oeschger events, may have contributed to the demise of Neanderthals is controversial. This type of variability has been pervasive during glacial periods (Margari et al., 2010; Martrat et al., 2004; Stockhecke et al., 2016) and Neanderthals, who moved into Eurasia ∼400 ka ago, had time to adapt to the related environmental changes. One can even make the argument that this adaptation to rapid temperature changes in Europe gave Neanderthals a strong advantage over AMHs, according to the variability selection paradigm (Potts, 1998). Dansgaard-Oeschger events are characterized by rapid warming and slow cooling (Barker et al., 2011; Martrat et al., 2004; Rasmussen et al., 2006; Thomas et al., 2009) (Fig. 4). A rapid opening of new climatically suitable corridors may have been beneficial for the dispersal of HS hunters and gatherers in Eurasia. The gradual cooling may have given them enough time to retreat southward to other ecological niches. To test this scenario, I conduct a series of simulations (**DOS**, Table 1), in which the level of Dansgaard-Oeschger variability in net primary production and temperature is varied from Φ=0,1,2 (Fig. 2d, 10).

Otherwise the simulations use the demographic parameters from the standard run **STD**. For the given standard parameters of interbreeding, competition, fecundity, diffusivity in **STD**, I find only small effects (<+/− 15%) of Dansgaard-Oeschger variability on the global population size of Neanderthals and *Homo sapiens* (Fig. 2d). Regionally, however, the effect of DO variability can be much more pronounced (**Supplementary Movies 1-4,** Figs. 5, 10). In Central Europe, DO events lead to an earlier Neanderthal extinction by about 500-1,500 years. In northern Europe, in particular over the Baltic Sea and northern Russia, the effect of DO events is much more proncouned. DO stadials can wipe out small populations which live at the fringes of their habitability range. Even though the subsequent return to DO warm events may create again favorable living conditions, the development of new HN founder populations in these areas is suppressed by the strong existing competition with *Homo sapiens* during this time. This effect can lead to a much earlier (up to ∼10,000 years) extinction of Neanderthals regionally. Averaged globally, in the absence of DO variability one finds an overall delayed extinction of Neanderthals of about 2,000-3,000 years, (Fig. 2d).

## 4. Summary and Discussion

This study set out to quantify the effects of competition, interbreeding and abrupt past climate change on the extinction of Neanderthals using a numerical 2-dimensional hominin dispersal model with realistic climate forcing and a set of initial parameters that mimic observed genetic and demographic asymmetries. The model simulations show that for low interbreeding rates, only competition can cause HN extinction at realistic times. Competitive exclusion in HDM2 is a consequence of the assumption that HS are twice as successful in using existing food resources compared to HN (K_S_/K_N_>2) (Neuhauser and Pacala, 1999). Moreover, in a time-varying environment HS gain further advantages over HN due to higher growth rates and resulting faster migration speeds *u*_S_. This allows HS to re-populate areas more quickly that underwent a rapid decline in NPP, for instance after DO stadial events. In such areas, higher initial HS densities lead to disadvantageous conditions for HN through the competition term. Parameter uncertainties were thoroughly tested in the presented parameter ensemble sensitivity experiments (**ENS, DOS, IBR**) to test the robustness of the major conclusions and develop a better understanding about the underlying processes, as well as their potential synergies.

Summarizing, this study confirms the notion that innovative generalists (HS) with higher levels of mobility, fecundity and plasticity, outcompete specialists (HN), who lived in smaller and more fragmented habitats (Melchionna et al., 2018). What specifically constituted the key competitive advantage of HS over HN is difficult to ascertain. It may have been a combination of different factors, including technological innovation of the Aurignacian industry of the Upper Paleolithic, with more wide-spread use of blades and bone and antler tools - mostly attributed to HS - over the Mousterian toolkits of the Middle Paleolithic, which were widely used by HN. Moreover, more sophisticated hunting techniques with possible use of domesticated dogs (Ovodov et al., 2011) as well as stronger resistance to pathogens originating from the African continent (Houldcroft and Underdown, 2016) may have given HS important longterm advantages, which would have contributed to the gradual competitive exclusion of HN.

The results presented here show that abrupt climate change, even though important regionally (Fig. 10), likely played only a minor role in the fate of the global Neanderthal population. According to the model simulations, interbreeding and assimilation, which have been proposed as important factors in the demise of European Neanderthal populations (Trinkaus, 2007) are effective only for low levels of food competition. To further constrain the effect of interbreeding in my model, more accurate genetic estimates of the total number of interbreeding individuals over the last glacial period would be necessary (Fig. 6a). However, currently, these numbers are still highly uncertain.

The Hominin Dispersal Model, version 2 (HDM2) provides a numerical framework to test existing hypotheses for Neanderthal extinction in terms of sensitivities with respect to bulk parameters of mobility, growth rate, competitiveness and interbreeding. More specific ideas, for instance on social or cultural superiority of HS (Gilpin et al., 2016) over HN, or on differences in anatomy (Hora and Sladek, 2014), hunting techniques or resistance to pathogens (Greenbaum et al., 2019) need to be translated into the bulk parameters of the phenomenological mean field model for hominin density. Another alternative approach to simulate such effects more explicitly is through agent-based modeling (Vandati et al., 2019). This is clearly an interesting area for future research.

The simulations presented here show that interbreeding between HN and HS is a relatively complex spatio-temporal phenomenon (Fig. 9), which starts in Anatolia and spreads westwards into the Balkans and reaches its peak in central Europe around 43 ka. It should be noted that earlier migrations of HS out of Africa into the Levant region (Timmermann and Friedrich, 2016) and potential encounters with HN there may have already contributed to the DNA exchange and low levels of admixture between these two groups. To simulate the effect of multiple interbreeding episodes in the model, one needs to choose earlier initial conditions (∼125 ka). This will be investigated further in future studies. My simulations of interbreeding and hybridization of populations (Fig. 9), provide additional information that can be further tested using genomic datasets from the last ice age (Fu et al., 2016).

The HDM2 is the first spatio-temporal model to simulate Neanderthal extinction in a temporally varying climatic environment. Whenever possible, model parameters were estimated based on previous genetic or demographic constraints. This has helped to capture in the model simulations key features of the dispersal, and interaction of the two hominin groups. However, there are still discrepancies between model experiments and archeological data (Fig. 8), namely the simulated presence of a HN population west of the Ural, which is absent in the archeological record. I hope that as new fossil, genetic and archeological data become available in future, key parameters in HDM2 might be further constrained, leading eventually to more realistic simulations of Neanderthal extinction across wider swaths of Eurasia.

### CRediT authorshop contribution statement

Axel Timmermann: Conceptionalization, Methodology, Investigation, Validation, Writing – review & editing, Software, Data curation.

### Data and Code availability

The HDM2 matlab code and all netcdf forcings files are available on: https://climatedata.ibs.re.kr/data/papers/timmermann-2020-QSR

## Acknowledgements

This work was supported by Institute for Basic Science (IBS) under IBS-R028-D1 and the Pusan National University Research Grant, 2017. Simulations were conducted on the IBS/ICCP Cray XC-50 supercomputer Aleph.

## Appendix A: Supplementary Data

**Supplementary movie 1.** Simulated HN population size (individuals/km^2^) for Standard simulation (STD) with realistic interbreeding and competition, which is initialized in 100 ka. Time counter counts forward in steps of 100 years starting from year 100 ka (counter 1).

**Supplementary movie 2.** Simulated HS population size (individuals/km^2^) for Standard simulation (STD) with realistic interbreeding and competition, which is initialized in 100 ka. Time counter counts forward in steps of 100 years starting from year 100 ka (counter 1).

**Supplementary movie 3.** Simulated HN population size (individuals/km^2^) for Baseline Simulation (BSL) without interbreeding and competition, which is initialized in 100 ka. Time counter counts forward in steps of 100 years starting from year 100 ka (counter 1).

**Supplementary movie 4.** Simulated interbreeding term (individuals/year/km^2^) for Standard Simulation (STD) with realistic interbreeding and competition, which is initialized in 100 ka. Time counter counts forward in steps of 100 years starting from year 80 ka (counter 300). Negative (positive) values correspond to the situation where hybrid offspring, originating from HN and HS interbreeding, assimilates into the HS (HN) group, (equation 1 Methods).

